# Sugar feeding patterns of New York *Aedes albopictus* mosquitoes are affected by environmental dryness, flowers, and host seeking

**DOI:** 10.1101/2020.03.26.009779

**Authors:** Kara Fikrig, Sonile Peck, Peter Deckerman, Sharon Dang, Kimberly St Fleur, Henry Goldsmith, Sophia Qu, Hannah Rosenthal, Laura C. Harrington

## Abstract

**Background:** Sugar feeding is an important behavior which may determine vector potential of mosquitoes. Sugar meals can reduce blood feeding frequency, enhance survival, and decrease fecundity, as well as provide energetic reserves to fuel energy intensive behaviors such as mating and host seeking. Sugar feeding behavior also can be harnessed for vector control (e.g. attractive toxic sugar baits). Few studies have addressed sugar feeding of *Aedes albopictus*, a vector of arboviruses of public health importance, including dengue and Zika viruses. To address this knowledge gap, we assessed sugar feeding patterns of *Ae. albopictus* for the first time in its invasive northeastern USA range.

**Methodology/ Principal Findings:** Using the cold anthrone fructose assay with robust sample sizes, we demonstrated that a large percentage of both male (49.6%) and female (41.8%) *Ae. albopictus* fed on plant or homopteran derived sugar sources within 24 hrs of capture. Our results suggest that sugar feeding behavior increases when environmental conditions are dry and may vary by behavioral status (host seeking vs. resting). Furthermore, mosquitoes collected on properties with flowers (>3 blooms) had higher fructose concentrations compared to those collected from properties with few to no flowers (0-3).

**Conclusions/Significance:** Our results provide the first evidence of *Ae. albopictus* sugar feeding behavior in the Northeastern US and reveal relatively high rates of sugar feeding. These results suggest the potential success for regional deployment of toxic sugar baits. In addition, we demonstrate the impact of several environmental and mosquito parameters (environmental dryness, presence of flowers, host seeking status, and sex) on sugar feeding. Placing sugar feeding behavior in the context of these environmental and mosquito parameters provides further insight into spatiotemporal dynamics of feeding behavior for *Ae. albopictus*, and in turn, provides information for evidence-based control decisions.

## Introduction

*Ae. albopictus* is a vector of numerous pathogens, including dengue, Chikungunya and Zika viruses as well as dog heartworm parasites [1, 2]. It is rapidly expanding its global range and is pushing northward in the USA, enabled by local adaptation and winter egg diapause [3, 4]. This highly adaptable mosquito can survive in drastically varied ecosystems, ranging from tropical to temperate climates, making it one of the most successful invasive species globally [5]. Understanding this mosquito’s feeding behavior and ecology across its invasive range is essential for understanding risk and devising control methods.

Sugar feeding is an important mosquito behavior with implications for disease transmission and control [6]. It can impact mosquito life history through a number of mechanisms and can vary between mosquito species [7]. For females, there may be trade-offs in transmission potential between blood and sugar feeding [7], as the latter may lead to satiation and reduce available abdominal space for a blood meal shortly after sugar feeding [8]. Sugar feeding has been considered a blood feeding suppressant in *Ae. albopictus* and other mosquito species, reducing blood meal size and frequency, thereby reducing opportunities for pathogen transmission [7, 9, 10]. In contrast, sugar feeding can enhance survival of *Ae. albopictus* and other mosquito species in laboratory studies, potentially increasing pathogen transmission [7, 9, 11-16]. Sugar also may enhance male mosquito mating performance by providing energy for mate-seeking and swarming [12, 16-19] and enable female host-seeking behavior [19]. In addition to impacts on mosquito life history, sugar feeding behavior has implications for the success of certain control and surveillance methods, such as attractive toxic (or targeted) sugar baits (ATSBs), which contain sugar and flower-derived attractants mixed with insecticides [20].

Environmental drivers that vary widely across *Ae. albopictus* invasive range can influence feeding behavior. For example, fructose feeding rates can vary by season and location, which may be caused by differences in temperature and humidity [21-25]. Dryness may stimulate sugar feeding behavior as has been shown by Hagan et al. (2018) for blood feeding [26]. Availability of sugar sources such as floral nectar may also affect mosquito sugar feeding rates, especially in arid climates [23, 27-29]. However, in addition to flower nectar, mosquitoes can acquire sugar from plant leaves, fruit, and homopteran honeydew [14, 23, 24, 30, 31] and these alternate sources can vary across *Ae. albopictus* habitats.

Given the importance of sugar feeding for mosquito fitness and the public health threat of *Ae. albopictus*, we know surprisingly little about its sugar feeding patterns in nature. Only four field studies have been conducted to date across vastly different habitats [16, 23, 32, 33]. Two studies indicated that season, habitat, and sugar availability might be important, as well as temperature and humidity [16, 23].

In Israel, the percent of sugar positive *Ae. albopictus* varied by season and habitat type (irrigated garden versus dry wasteland), ranging from 41.4% (summer) to 74.1% (fall) at a wasteland site [23]. However, this study was performed with an abbreviated cold anthrone assay, using visual detection of color change, instead of precise measurement of fructose concentration using established methods [23]. A subsequent study evaluating *Ae. albopictus* visitation to sugar sources reported attraction to a subset of tested ornamental flowers, wildflowers, damaged carob seed pods and fruits, but no attraction was detected to honeydew coated plants [33]. Working with releases of high generation laboratory colony males in northern Italy, Bellini et al. (2014) utilized an abbreviated cold anthrone assay to detect higher sugar feeding rates for released males at sites with sucrose feeding stations compared to control sites and a positive correlation with temperature and negative correlation with humidity [16]. In Florida, where the only other US study was conducted, *Ae. albopictus* fructose concentration did not vary significantly with plant species utilized as resting habitat; unfortunately, no analyses were conducted to determine the proportion sugar fed [32]. Another limitation of these studies was the lack of established baseline fructose levels, leading to the potential misidentification of larval nutrients as adult sugar meals.

The current knowledge of *Ae. albopictus* sugar feeding in the field primarily stems from these four studies in Israel, Italy, and Florida. Additionally, assessments of ATSBs for *Ae. albopictus* population control in Florida and Israel have demonstrated that sugar feeding frequency is sufficient to achieve population reductions [34-38]. However, these locations are not representative of the vast environmental variation in climate and flora where *Ae. albopictus* is now established. It has yet to be determined whether sugar feeding behavior of *Ae. albopictus* in other regions of the world will be conducive to ATSB success due to an absence of basic ecological and behavioral information on this subject.

To address this important gap, we assessed the sugar feeding behavior of *Ae. albopictus* at its invasive edge in Northeastern USA in order to understand its feeding ecology along the northern limit of its expanding range. We determined the proportion of male and female mosquitoes that contained fructose and individual mosquito fructose concentrations. In addition, we assessed the response of sugar feeding to environmental dryness, floral presence, host seeking status, and sex. Placing sugar feeding behavior in the context of these environmental and mosquito parameters provides further insight into spatiotemporal dynamics of this behavior for *Ae. albopictus*, and in turn, provides information for evidence-based control decisions.

## Methods

### Field Site

Mosquitoes were collected in Long Island, New York, USA, at four farms and four residential areas with between nine and seventeen houses in each, totaling fifty properties. Sites were chosen based on prior knowledge of *Ae. albopictus* distribution in Suffolk County from larval surveys and vector control surveillance [39](S. Campbell, pers comm). Collections were conducted between June and August 2018. Two HOBO Pro v2 data loggers (model U23-001, Onset Computer Corp., Bourne MA, USA) per site recorded the temperature and humidity every four hours from mid-July through August.

### Mosquitoes

Resting mosquitoes were collected using large custom-designed aspirators (30.5 cm diameter, 114 cm height, 12 V PM DC 2350 RPM, 1/35 Horse power, 3.7 Amp motor) [40] and host seeking mosquitoes that approached collectors were caught with nets. Collection bags were placed in acetone jars for ∼3 min. Anesthetized mosquitoes were then separated into microcentrifuge tubes, placed on ice and transported to the laboratory. Mosquitoes were identified to species using published keys [41], sorted by blood meal status, labeled, and stored at -20°C. A small number of the blood-fed mosquitoes (182) were saved for blood meal analysis (Fikrig, unpublished data) and non-blood fed mosquitoes were utilized for sugar analysis. Mosquitoes were transported on dry ice to Cornell University for further processing. To determine body size, one wing was removed from each mosquito, placed on a slide and measured from the axillary incision to base of fringe hairs [42] using a dissecting microscope and software (Olympus SZX9, Olympus DP22 camera, and Olympus cellSens software).

### Flower Census

Beginning in mid-July 2018, the number of blooming flowers per morphospecies (up to 100 blooms) was counted on each property (n=54). Morphospecies (morphologically distinct species) were identified using the GardenAnswers phone application (Garden Answers LLC., San Diego, CA) [43]. Flowers were categorized into groups representing flower presence: absent (0-3 blooms) and present (>3 blooms). A range from zero to three was chosen to represent an absence of flowers rather than zero because three was a natural breaking point in the data, with only one mosquito collected on a property with 6 or 8 flowers, and all others on properties with at least 9 flowers, creating a natural gap between mosquitoes collected on properties with 0-3 and the rest of the dataset. Properties with up to three flowers had relatively little nectar and were therefore considered an appropriate comparison group to more highly flowered properties, expanding the number of mosquito observations on ‘absent’ properties by 50% compared to an absolute zero ‘absent’ group.

### Fructose Detection

#### Cold Anthrone Assay

Fructose concentration was measured using the cold anthrone colorimetric assay [44]. At room temperature, anthrone solution reacts with fructose, but not other sugars. The assay is indicative of plant feeding and does not measure blood sugars (primarily glucose) or stored sugars (trehalose), although non-sugar fed teneral mosquitoes contain small amounts of fructose.

Mosquitoes were homogenized in 1.7 ml microcentrifuge tubes using a lyser (FastPrep-24™ Classic Instrument, MP Biomedicals, USA) at 4 m/s for 30 s with 50 µl of 2% sodium sulfate solution and glass beads (3 mm, Thermofisher). Chloroform methanol (1:2) solution (375 µl) was added to each tube and vortexed for 8 s and centrifuged for 15 min at 200 x g, extracting fructose into supernatant. Tubes were stored at -20°C until fructose quantification, at which time 10 µl of supernatant was transferred to two wells of a 96-well microplate.

To ensure consistency, standards were created once via serial dilution and stored at 4°C for the duration of analysis. Two replicates of standards (10 µl of 0, 0.05, 0.1 and 0.2 µg /µl D-Fructose (Fisher Chemical, USA) in 25% ethanol) and samples (10 µl) were pipetted individually into wells on each plate. Thereafter, 240 µl anthrone solution (freshly prepared each day, 67.89 µl distilled water, 172.1 µl sulfuric acid, and 0.339 mg anthrone per sample) was added with a multichannel pipette. Samples were incubated at room temperature for 90 min in a chemical hood. The absorbance of light (630 nm) by the reaction of each sample was measured by the microplate reader (800 TS Absorbance Reader, BioTek, VT, USA) and compared against the standard curve to determine fructose concentration. The mean of the two experimental replicates of each sample was used in analyses, except when experimental replicates were dissimilar, the data were discarded or sample was reanalyzed.

#### Baseline Mosquito Fructose Concentrations

During the period of adult collection, water-holding containers were examined from a subset of properties and pupae were collected and held in the laboratory until eclosion. Post-eclosion, adult *Ae. albopictus* (n= 77 male, 52 female) were held without sugar and frozen within 12 hrs of emergence. These mosquitoes were used to establish a field baseline level of fructose in teneral *Ae. albopictus* [21]. Adults with fructose concentrations greater than one standard deviation above the sex-specific mean baseline concentration were considered to be fructose-positive.

#### Laboratory Digestion Assay

To determine the time window of fructose detection in *Ae. albopictus* post-sugar meal consumption, we conducted an assay of fructose concentration in sugar fed females over digestion time [45]. *Ae. albopictus* (F6 from NY at 24°C and F8 from FL at 28°C) were fed 10% sucrose solution between 1 – 3 d post-eclosion. Males and females were removed before, immediately after, and at 24-hour intervals post sugar feeding. Between nine and twenty mosquitoes were removed per day. Fructose concentration was measured as described above. The assumption of constant variance was not met, so mean fructose concentrations were compared to concentration before feeding using the non-parametric Kruskal-Wallis test followed by a Dunn’s multiple comparisons test with Benjamini-Hochberg correction (reported as P_adj_).

## Data Analysis of Sugar Feeding Patterns in the Field

Analyses were performed in R (Version 1.1.463) [46]. Average fructose concentration and proportion sugar fed were calculated for all male and female *Ae. albopictus* collected from June to August 2018. Wing measurements were used to standardize fructose concentration by body size, by dividing total concentration by µm wing length. A subset of *Ae. albopictus* for which we had flower and weather data (those collected between 23 July - 15 August) were included in the models described hereafter.

A Generalized Linear Mixed Model (GLMM; lme4 package) with binomial distribution was employed to determine the impact of measured variables on sugar feeding probability of a captured mosquito [47]. Random effects included town, address nested in town, date, town-date interaction, and address nested in town-date. Fixed effects included capture method (aspirator or net), sex, presence of open flower blooms on property, and saturation deficit. Initially, several weather parameters were evaluated, including minimum, maximum, and average temperature and humidity, as well as saturation deficit [48].

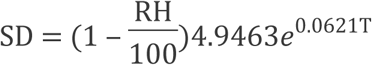

All weather parameters had similar explanatory power in the models, so saturation deficit was chosen for the final model because it included both temperature and humidity in a biologically relevant way. Because the sugar was detectable for up to 24 hrs after consumption in our laboratory assessments, the cumulative saturation deficit over that time was determined by summing the saturation deficit over the 6 most recent time points (a 24 hr interval) prior to collection time for each mosquito. Flower count was included as a binary variable measuring flower presence as described above.

A linear mixed model was employed to evaluate log fructose concentration standardized by wing length using all the fixed and random effects listed above. Only mosquitoes that were sugar fed were included in this analysis to further understand factors influencing the magnitude of sugar feeding.

For both GLMM and linear mixed models, the emmeans package was used to conduct post-hoc analyses of the effects of individual parameters [49].

## Results

### Environmental and flower measurements

For the dates July 23 – August 15, 2018, mean ± SD temperature was 24.4 ± 2.79°C (range 15.4°C - 37.1°C). Mean relative humidity was 87.1 ± 12.2 % (range 1%-100%). Floral counts varied by property visit (collection event on a given property): more properties had flowers present (154) during mosquito collections than absent (20). The median number of flowers per property was 110.5.

### Mosquitoes

Between June and August 2018, 2,788 *Ae. albopictus* were collected, representing 1,263 females (45.3%) and 1,525 males (54.7%). Of these, 2,517 (90.3%) were collected resting on vegetation and other surfaces by aspirator and 271 (9.7%) were captured flying around human collectors with nets (241 female and 30 male). Mosquitoes were captured across 8 sites, with 1,097 (39.3%) from the four farms and 1,691 (60.7%) from the four residential areas. Among the subset of mosquitoes that were captured during floral census (1,970), more were collected on properties with flowers present (1,827, 92.7%) than flowers absent (143, 7.26%).

Females wings were 2.71 ± 0.27µm (mean ±SD; range: 1.59 – 3.50µm) and male wings were 2.22 ± 0.22µm (range 1.24 – 3.21µm).

### Fructose Detection

#### Field-caught mosquitoes

Among mosquitoes collected from June through August, a relatively high proportion of both male (756/1,525, 49.6%) and female (528/1,263, 41.8%) *Ae. albopictus* were sugar fed. The percent of sugar fed mosquitoes by each variable is displayed in Table 1. Among sugar fed mosquitos, average female fructose concentration was 0.0488 µg/µl and male fructose concentration was 0.0300 µg/µl. To account for differences in body size, fructose concentrations were standardized by wing length for sugar fed females (0.0180 ± 0.0182 µg/(µl*µm)) and males (0.0134 ± 0.0132 µg/(µl*µm)).

**Table 1.**
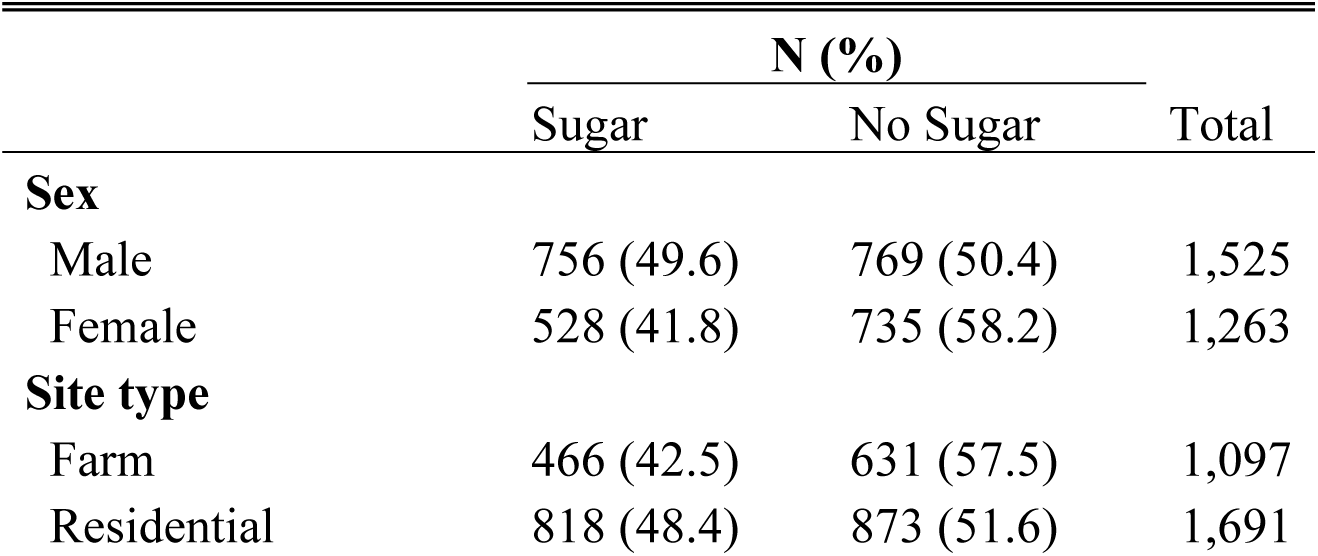
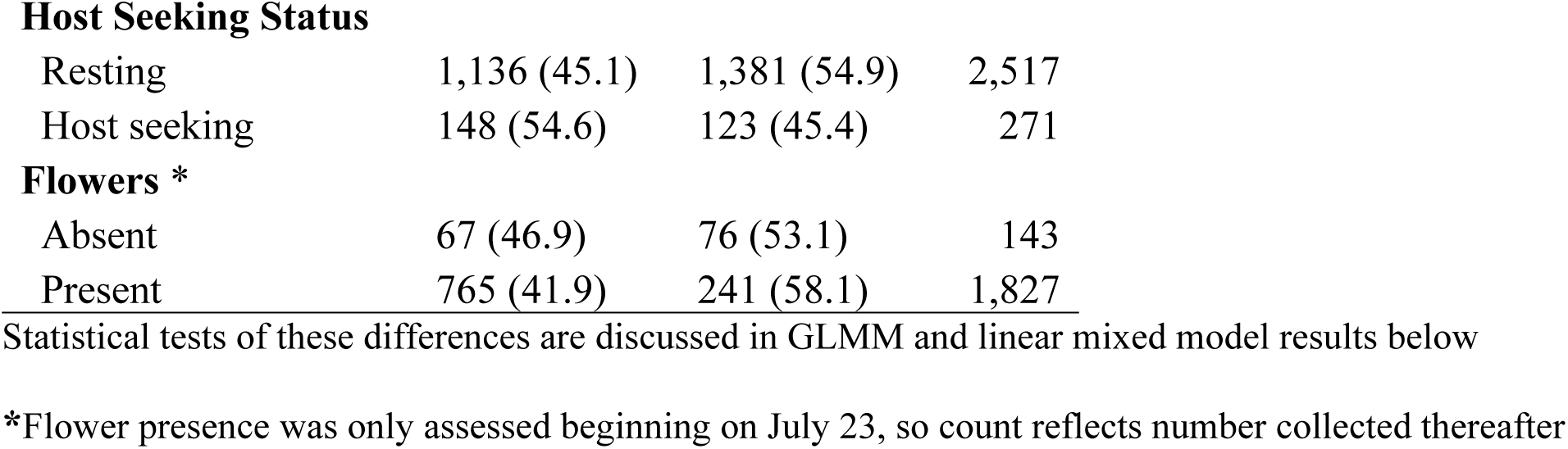
Sugar fed status of mosquitoes captured by site, status, sex and floral abundance.

|  | N (%) |  | Total |
| --- | --- | --- | --- |
|  | Sugar | No Sugar |  |
| **Sex** |  |  |  |
| Male | 756 (49.6) | 769 (50.4) | 1,525 |
| Female | 528 (41.8) | 735 (58.2) | 1,263 |
| **Site type** |  |  |  |
| Farm | 466 (42.5) | 631 (57.5) | 1,097 |
| Residential | 818 (48.4) | 873 (51.6) | 1,691 |
| **Host Seeking Status** |  |  |  |
| Resting | 1,136 (45.1) | 1,381 (54.9) | 2,517 |
| Host seeking | 148 (54.6) | 123 (45.4) | 271 |
| **Flowers \*** |  |  |  |
| Absent | 67 (46.9) | 76 (53.1) | 143 |
| Present | 765 (41.9) | 241 (58.1) | 1,827 |

Statistical tests of these differences are discussed in GLMM and linear mixed model results below
\*Flower presence was only assessed beginning on July 23, so count reflects number collected thereafter

#### Baseline Fructose Concentration

Mean ± SD fructose concentrations from field collected pupae that eclosed in the laboratory without sugar were 1.42 ± 2.76 ng/µl for females and 0.935 ± 2.09 ng/µl for males. Baseline concentrations (mean fructose concentration +1 SD) were 4.18 ng/µl for females and 3.02 ng/µl for males. All field-captured adult fructose values above this baseline level were considered sugar fed.

#### Laboratory Digestion Assay

Male and female *Ae. albopictus* digested fructose within 24 hrs after ingestion (Fig 1) at both low (23.5°C) and high (28°C) constant temperatures. Compared to fructose levels before sugar feeding (Day 0), fructose was only detectable on Day 1, immediately after feeding (Kruskal-Wallis test with post-hoc Dunn’s test; female 23.5°C: P_adj_=0.0051; male 23.5°C: P_adj_=0.0004; female 28°C: P=0.0099; male 28°C: P=0.0001). At the next check point, 24 hrs post-feeding (Day 2), and all days thereafter (Days 3-6), fructose concentrations were either not different from (Kruskal-Wallis post-hoc Dunn’s, P_adj_>0.05) or lower (Female 28°C Day 5 : P_adj_=0.0105 and Day 6: P_adj_=0.0052) than concentrations before sugar feeding.

**Fig 1:**
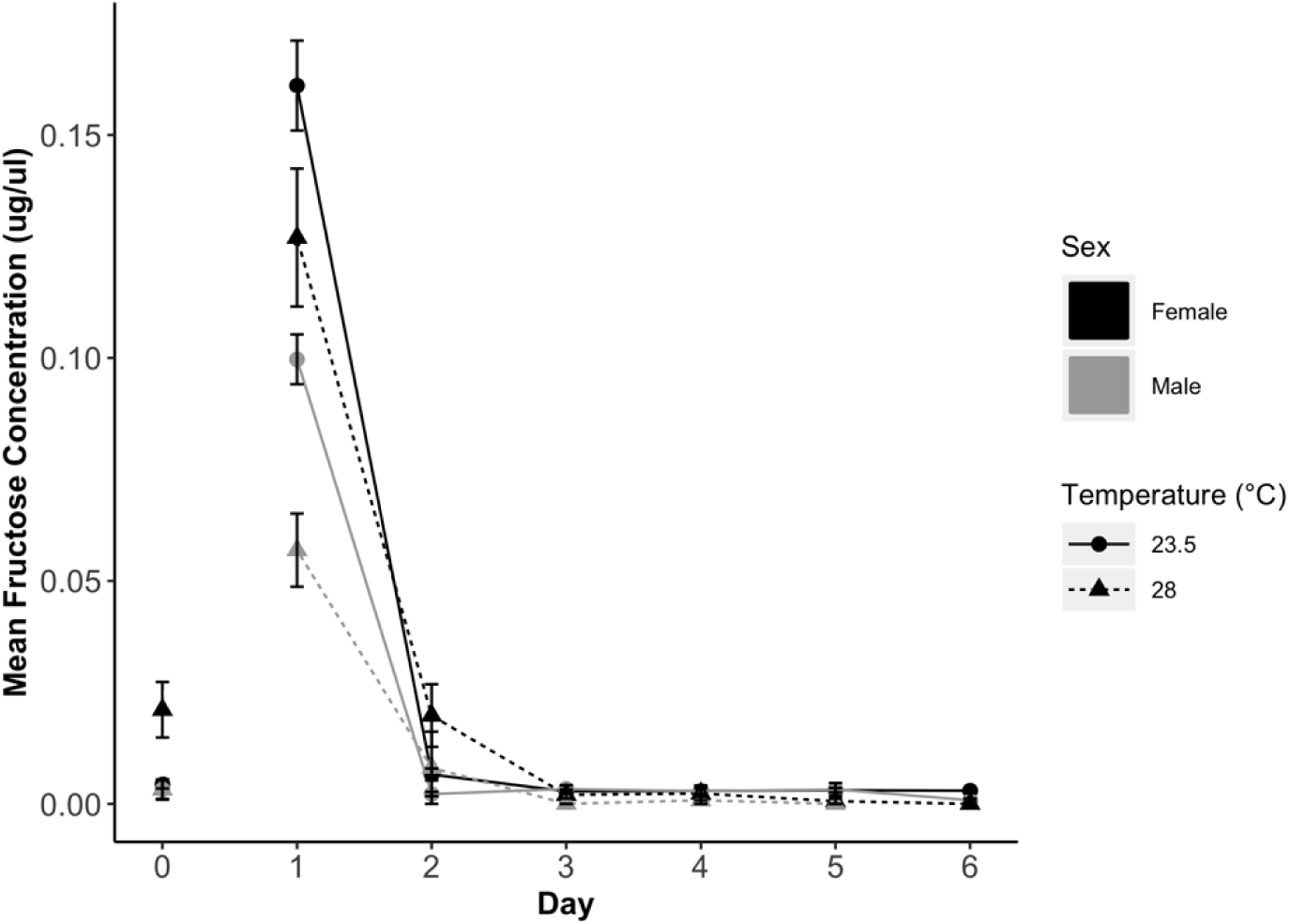
Male and female digestion of fructose over time at 23.5C and 28C. Fructose concentration was measured daily after time of ingestion (Day 1). The daily mean (±SE) fructose concentration is shown for females (black) and males (gray) at 23.5°C (solid line and circle points) and 28°C (dotted line and triangle points). Compared to pre-ingestion fructose concentrations (Day 0), mosquitoes only had significantly higher fructose concentrations on Day 1 (immediately after ingestion) for each sex at each temperature (Kruskal-Wallis Dunn’s test; P_adj_<0.05).

### Adult *Ae. albopictus* sugar feeding patterns

#### Effects of environmental and mosquito parameters on sugar feeding status

For the subset of mosquitoes captured after floral and weather data collection was initiated, saturation deficit, host seeking status, and sex influenced the probability of sugar feeding while the number of flowers on a property did not (GLMM, P value). The likelihood of sugar feeding was affected by dryness as measured by saturation deficit (n=1,970, β=0.0470, SE=0.0148, *P*=0.00143). More mosquitoes fed on sugar when the saturation deficit was high (i.e. when weather was hotter and drier) (Figure 2). Host seeking mosquitoes (n=151) were more likely to be sugar fed than resting individuals (n=1,673; β=0.527, SE=0.201, *P*=0.00870). Males (n=1,042) were more likely to be sugar fed compared to females (n=782; β=0.394, SE=0.110, *P*=0.000321). The relative abundance of flowers did not affect the likelihood of sugar feeding; mosquitoes collected on properties with flowers (n=1,827) were not more likely to be sugar fed than those captured on properties with no flowers (n=143; β= 0.0728, SE=0.290, *P*=0.802).

**Fig 2:**
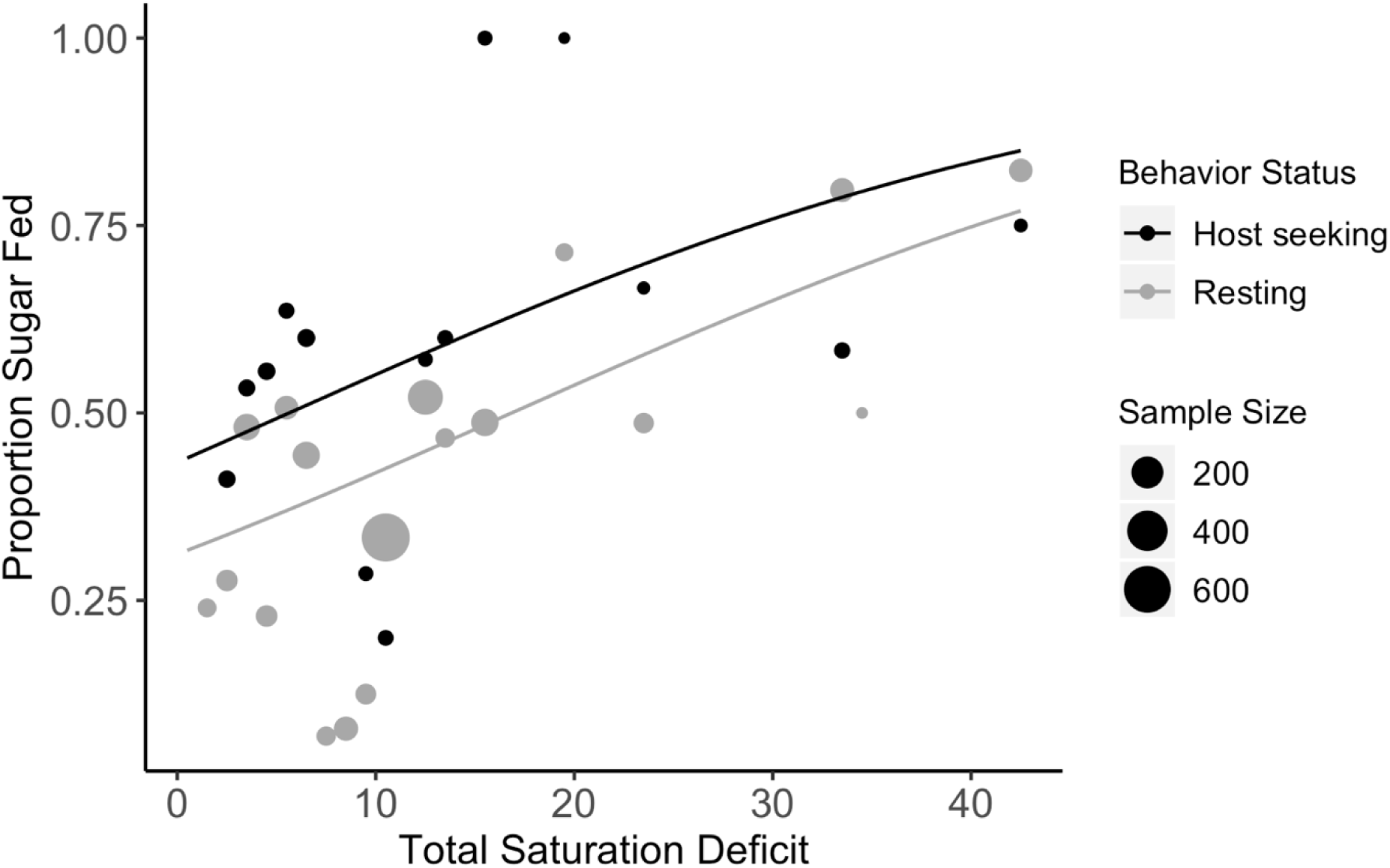
The proportion of sugar fed mosquitoes by saturation deficit for host seeking (black) and resting (gray) mosquitoes. Mosquitoes were grouped by 1 unit of saturation deficit. The total number of mosquitoes collected per unit saturation deficit is represented by point size. The predicted probability of sugar feeding by saturation deficit is indicated by the lines. As saturation deficit increased, the likelihood of capturing a sugar-fed mosquito increased (GLMM, *P*=0.00143). Mosquitoes captured while host seeking were more likely to be sugar fed than while resting (GLMM, *P*=0.00870) [49].

**Fig 3:**
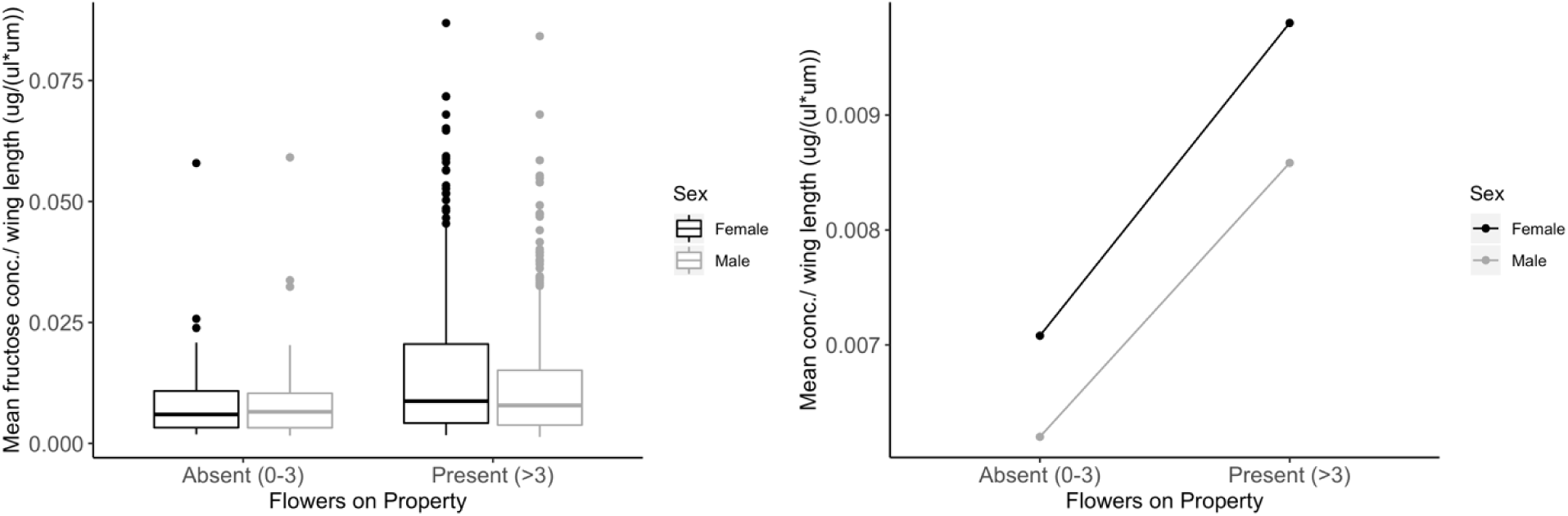
**A. Mean fructose concentration standardized by wing length for female (black) and male (gray) *Ae. albopictus* on properties with and without flowers.** Points show individual mosquito fructose concentration standardized by wing length of outliers. **B. Predicted fructose concentration by flower presence and sex**. Mosquitoes collected on properties with flowers present (LMM, *P*=0.0253) had higher fructose concentration per µm wing length than those collected on properties with flowers absent. Females had marginally higher fructose concentration compared to males (LMM, *P*=0.0553).

#### Effects of environmental and mosquito parameters on fructose concentration ingested

The linear mixed model results showed that the fructose concentration in sugar fed mosquitoes was predicted by flower abundance but not by environmental dryness, sex, or host seeking status. Among sugar fed mosquitoes (n=832), those collected on properties with flowers present (n=765) had significantly higher fructose concentration per µm wing length than those collected on properties with flowers absent (n=67) (β=0.325, SE=0.142, *P*=0.0253). Males (n=507) took marginally smaller sugar meals compared to females (n=325) even when controlling for body size differences between the sexes (β=-0.133, SE=0.0691, *P*=0.0553). There was no significant effect of host seeking status (host seeking vs resting, β=0.217, SE=0.116, *P*=0.0611) or saturation deficit (β=0.00844, SE=0.00711, *P*=0.253) on fructose concentration per µm wing length.

## Discussion

Sugar feeding patterns of field captured *Ae. albopictus* mosquitoes have only been reported from three other locations: Israel, Italy, and Florida [16, 23, 32]. Our study reports Asian tiger mosquito sugar feeding patterns for the first time from the northern edge of its invasion in the Eastern USA. Using robust sample sizes, we demonstrated that a large proportion of both male and female *Ae. albopictus* fed on plant or homopteran derived sugar sources within 24 hrs of capture. Our results suggest that sugar feeding behavior increases when environmental conditions are dry and may vary by behavioral status (host seeking vs resting). Furthermore, mosquitoes collected on properties with flowers had higher fructose concentrations compared to those collected from properties with no flowers.

A large percentage of males (49.6%) and females (41.8%) collected from our field sites were sugar fed. Our laboratory assays demonstrated that *Ae. albopictus* digest fructose within 24 hrs of consuming a sugar meal at 23.5 and 28°C. According to this window of detection, and considering the average field temperatures during collections, approximately half of the field-captured mosquitoes fed on sugar daily. Sugar feeding estimates may be influenced by the concentration of sugar consumed, which varies between flower species’ nectar and between alternative sources of sugar. In our study we used a 10%-sucrose solution representing the low end of sugar concentrations in nectar (7-70%)[50]. In a temperate region of Italy, similar rates of sugar feeding were detected among released males (48% at 72 hours post-release) compared with wild males in our study[16]. In the arid climate of Israel, sugar feeding tended to be more common; the percentage of sugar fed mosquitoes ranged from 41.3% to 74.1% based on season and site [23].

In Long Island, *Ae. albopictus* that experienced higher saturation deficits (hotter, drier weather) 24 hours prior to collection were more likely to contain a sugar meal than those collected during lower saturation deficits. Bellini et al. (2014) observed a similar pattern with field-released males when assessing sugar feeding devices; the percentage of sugar positive males was correlated negatively with relative humidity and positively with temperature at control sites [16]. It is possible that high saturation deficit leads to dehydration and ultimately triggers higher rates of sugar feeding, especially on more dilute sources. Maintaining water balance is essential for insect survival [51, 52] and others have described insect foraging behaviors that balance physiological needs for water and sugar through choice of nectar dilution levels [53-56]. Working with mosquitoes, Hagan et al. (2018) found that blood feeding was prompted by dehydration. Although sugar and blood feeding are different behaviors and dilute nectars can contain similar or lower levels of water compared to blood, it is possible that mosquitoes use the same set of physiological cues to prompt sugar feeding under dehydrating conditions [26].

In our study, the presence of flowers did not influence the likelihood of *Ae. albopictus* sugar feeding but did impact the amount of sugar ingested when they did feed. Yard sizes varied in our study but tended to be small and within the flight range of *Ae. albopictus* [57, 58], so it is possible that sugar fed mosquitoes collected in yards without flowers originally sugar fed in adjacent yards with greater floral abundance, and subsequently used some of the fructose in flight. This could explain why we observed consistent likelihood to sugar feed between flower categories, but different fructose concentrations between mosquitoes collected on properties with flowers present and absent. Alternatively, sugar fed mosquitoes in yards without flowers may have consumed non-nectar sources, such as honeydew or plant tissue. Parasitoid wasps fed on honeydew had lower fructose levels compared to those fed on nectar [59] and plant leaves generally have lower concentration of sugars than nectar [60]. This would also account for the equal likelihood of feeding and different fructose concentrations by flower presence.

Only one other published study has investigated floral abundance and *Ae. albopictus* sugar feeding and differs from our results. In Israel (under arid environmental conditions), a difference in sugar feeding likelihood was observed by flower abundance: 42% and 68% of females from low and high floral abundance sites, respectively, contained sugar [23]. It is difficult to compare our study with that of Müller et al. (2010) because of vastly different environmental conditions. Houses without flowers in Long Island still had significant vegetation and potential non-nectar sugar availability, in contrast to the less vegetated “dry wasteland” site in Israel. Sampling limitations in our study restricted floral surveys to properties where mosquitoes were collected, preventing inclusion of flowers in neighboring yards within *Ae. albopictus* flight range. Quality of floral resource was also not considered, such as nectar quantity or quality, which can be highly variable [61].

A subset of host seeking mosquitoes were captured with nets as they flew around human collectors. These mosquitoes were more likely to contain sugar meals than those collected with aspirators while resting on vegetation and other surfaces. While some studies have reported reduced blood feeding after sugar feeding [7, 9, 10], it is possible that teneral females seek sugar meals shortly after eclosion before blood feeding [8], explaining higher sugar content in host seeking females. This observation warrants further investigation. In addition, it highlights the importance of considering collection method biases when assessing sugar feeding prevalence and should be an important consideration when designing and analyzing sugar feeding study results.

These sugar feeding patterns will likely influence the success of sugar-based control techniques, such as ATSBs. While this control strategy has only been assessed for *Ae. albopictus* populations in Florida and Israel [20, 34-38], our results provide insight into the potential for deployment in our study region. In Israel, 62.7% of female *Ae. albopictus* were sugar fed at a natural garden site; meanwhile, ATSB deployment reduced biting pressure by 85% at another site under similar conditions [23, 36]. The comparatively lower percentage of sugar fed females in Long Island (41.8%) may result in weaker reductions in biting pressure in our study region, but the sugar feeding rates were nevertheless sufficient to warrant further investigation of ATSB-based control methods in Northeastern US. Our results suggest that control success in our region may be maximized if ATSBs are deployed during hot, dry conditions and in locations with fewer flowers and less competition. Furthermore, the tendency of *Ae. albopictus* to sugar feed prior to blood feeding may increase the public health impact of ATSBs by concentrating control pressure before the point of pathogen acquisition or transmission.

As the ability to detect DNA from mosquito plant meals improves, future studies could explore sugar feeding with greater resolution than the cold anthrone test affords. Next-generation sequencing has been employed to successfully identify plant meals of mosquitoes and other blood feeding Diptera [62, 63]. Additional studies of *Ae. albopictus* plant meal origin would be beneficial in ATSB lure design optimization. However, results of these analyses must be interpreted with caution as they may bias towards non-nectar sugar sources that are more likely to be detected via DNA-based analyses due to minimal DNA content of nectar.

Our results demonstrate, for the first time, sugar feeding patterns by temperate populations of *Ae. albopictus* in the United States. This is only the fourth field study on this important mosquito behavior and provides us with insights into conditions that might influence sugar feeding variation, including atmospheric dryness, flower presence, and host seeking. In light of the high frequency of sugar feeding in the study population, our results show promise for deployment of attractive toxic sugar baits for *Ae. albopictus* control in the region and provide insight into potential modifications of bait timing and placement to maximize success.

## Acknowledgements

We thank Dr. Dan Gilrein at Cornell Cooperative Extension, Moses Cucura, and Dr. Scott Campbell at Suffolk County Government for providing laboratory space and logistical support, Dr. Erika Mudrak for providing statistical expertise, Dr. Talya Shragai for assistance in selecting sites, and Sylvie Pitcher for support in the lab.

